# Contribution of neurons that express *fruitless* and *Clock* transcription factors to behavioral rhythms and courtship

**DOI:** 10.1101/2024.06.12.598537

**Authors:** Anthony Deluca, Brooke Bascom, Daniela A Key Planas, Matthew A Kocher, Marielise Torres, Michelle N Arbeitman

**Affiliations:** Department of Biomedical Sciences, College of Medicine, Florida State University, Tallahassee, FL 32306

**Author notes:** Co-first authors (Anthony Deluca, Brooke Bascom, Daniela A Key Planas, Matthew A Kocher). The authors declare no competing interests.

**Keywords:** Drosophila, courtship, reproduction, sex determination, circadian rhythms, connectome

## Abstract

Animals need to integrate information across neuronal networks that direct reproductive behaviors and circadian rhythms. In Drosophila, the master regulatory transcription factors that direct courtship behaviors and circadian rhythms are co-expressed in a small set of neurons. In this study we investigate the role of these neurons in both males and females. We find sex-differences in the number of these *fruitless* and *Clock* -expressing neurons (*fru* ∩ *Clk* neurons) that is regulated by male-specific Fru. We assign the *fru* ∩ *Clk* neurons to the electron microscopy connectome that provides high resolution structural information. We also discover sex-differences in the number of *fru*-expressing neurons that are post-synaptic targets of *Clk*-expressing neurons, with more post-synaptic targets in males. When *fru* ∩ *Clk* neurons are activated or silenced, males have a shorter period length. Activation of *fru* ∩ *Clk* neurons also changes the rate a courtship behavior is performed. We find that activation and silencing *fru* ∩ *Clk* neurons impacts the molecular clock in the sLNv master pacemaker neurons, in a cell-nonautonomous manner. These results reveal how neurons that subserve the two processes, reproduction and circadian rhythms, can impact behavioral outcomes in a sex-specific manner.

## Introduction

Reproductive behaviors and circadian rhythms are biologically linked. For example, the time-of-day influences when animals mate, and reproductive behaviors and mating in turn impact daily activity patterns and sleep. In *Drosophila melanogaster*, the neuronal networks that direct reproduction and circadian rhythms are largely distinct, but there are neurons predicted to function in, or link, both processes, given that they express the master regulatory transcription factors that underlie both reproduction and daily rhythms. A set of transcription factors encoded by *period (per)*, *timeless (tim), Clock (Clk)*, and *Cycle* (*Cyc*) are part of a regulatory feedback loop to direct circadian rhythms ^1^. The sex determination hierarchy gene *fruitless* encodes sex-specific transcription factors that are produced in neurons that underlie male and female reproductive behaviors ^2^. In our previous single-cell RNA-sequencing study directed at molecularly classifying the repertoires of *fruitless*-expressing neurons that direct reproductive behaviors ^3^, we identified a set of neurons in both sexes that also express the master clock genes (*fru* ∩ *Clk* neurons). Here we examine the role of *fru* ∩ *Clk* neurons in generating sex-differences in behavioral rhythms and determine if there are additional functions for reproductive behaviors.

The network of neurons directing circadian rhythms is comprised of ∼150 neurons in the brain that have been ascribed different functions. These neurons are named for their neuroanatomical positions. The *fru* ∩ *Clk* neurons are defined by molecular-genetic tools and include a subset of each of the following classes of clock neurons: LNds (dorsal-lateral neurons), DN1s (dorsal) and DN3s (dorsal) clock neurons. The LNds direct evening anticipation in light:dark conditions (LD) and are part of the set of “evening” or “E” cells. The DN1s are targets of the sLNv “morning” or “M” neurons. The sLNvs are pacemaker neurons that are critical for behavioral activity rhythms and produce pigment dispersing factor (PDF), a neuropeptide that keeps the clock neuronal network synchronized. The DN1s have been ascribed several functions, including activity promoting ^4^, sleep promoting ^5,6^, and coordinating the response to light and temperature ^7^. The DN3s are not as well studied, though do have a role in promoting sleep ^8^.

The potential for Drosophila reproductive behaviors is specified by the sex determination hierarchy ^9^. This hierarchy is comprised of a pre-mRNA splicing cascade that is a read-out of the number of X chromosomes. At the bottom of the hierarchy are two genes that encode pre-mRNAs that are spliced in a sex-specific manner that control all aspects of sexual differentiation in non-germline tissues (*doublesex* [*dsx*] and *fruitless* [*fru*]). Both *dsx* and *fru* direct reproductive behaviors through production of sex-specific transcription factors, with *fru* being expressed in a larger population of neurons. *dsx* also directs all aspects of somatic sexual differentiation outside of the nervous system. The *fru* transcripts produced from the *fru P1* promoter are the sex-specifically spliced *fru* transcript class, that produces male-specific transcription factors (Fru^M^). It is thought that no functional female-specific Fru product is produced. *fru P1*-expressing neurons are found in both sexes and have similar neuroanatomical positions and numbers ^10,11^. These *fru P1* neurons underlie reproductive behaviors in both sexes, based on several types of molecular-genetic perturbation studies ^12^.

Here, we examine *fru* ∩ *Clk* neurons that are molecularly defined using a genetic intersectional approach that relies on *fru P1* regulatory elements driving expression of *flippase* (*fru-FLP*) ^13^, the *Clk856-Gal4* transgene (*Clk-Gal4*) that drives expression in the core clock neurons of UAS-controlled transgenes ^14^, and a set of *UAS-FLP-out* cassette transgenes that have expression limited to neurons with overlapping *fru-FLP* and *Clk-Gal4* expression. We extend our neuroanatomical examination of sex differences in these neurons using a MultiColor FlpOut (MCFO) stochastic labeling approach, coupled with links to the Drosophila FlyWire connectome that is based on serial electron micrographic studies. Furthermore, we identify sex differences in the number of *fru P1* neurons that are targets of the core clock neural network, using a trans-synaptic labeling approach (trans-TANGO) ^15^. We found that either activating or silencing the *fru* ∩ *Clk* neurons results in a shorter circadian period and more activity in constant dark conditions (DD), but only in males. Activation and silencing of *fru* ∩ *Clk* neurons had modest impacts on sleep and overall activity in LD conditions. Male courtship behavior was impacted by *fru* ∩ *Clk* neurons activation, with more attempts to copulate per minute observed, linking the *fru*-expressing circadian neurons to reproductive activity. We found that the changes in behavioral rhythmicity were reflected in Per cycling in the sLNvs neurons, when *fru* ∩ *Clk* neurons were either activated or silenced. Taken together, we have found links between sex-specific circadian governance of daily activity and reproductive outputs.

## Results

### Characterization of *fru* ∩ *Clk* neurons

To visualize neurons that express both *fru P1* (hereafter *fru*) and *Clk* (*fru* ∩ *Clk*) we employed a labeling approach called MultiColor FlpOut (MCFO) that relies on expression of both *fru-FLP* and *Clk-Gal4* for the visualization of individual neurons and their projections in different colors, using confocal microscopy (**Figure 1A**; called *fru* ∩ *Clk* MCFO neurons). ^13,14,16^. We first observed *fru* ∩ *Clk* MCFO neurons over a 24-hour period, in animals that are aged 5-10 days old, across 6 time points, to determine if there were changes in neuronal projection patterns influenced by the time of day, as observed for sLNvs ^17^. We did not observe any robust changes in neuronal projection patterns (**Supplemental Figure 1**). We did an in-depth characterization of the full set of *fru* ∩ *Clk* MCFO neurons from animals at Zeitgeber Time (ZT) 4-6. We observe expression in a subset of the DN1s, DN3s and LNds, as before ^3^, with projections within the protocerebral region for the DN1s and DN3s, including some with projections that cross the midline. Additionally, our labeling identifies neurons in the subesophageal zone (SEZ), ascending median bundle neurons and in the ventral nerve cord (VNC), with most in the abdominal ganglion of the VNC.

**Figure 1.**
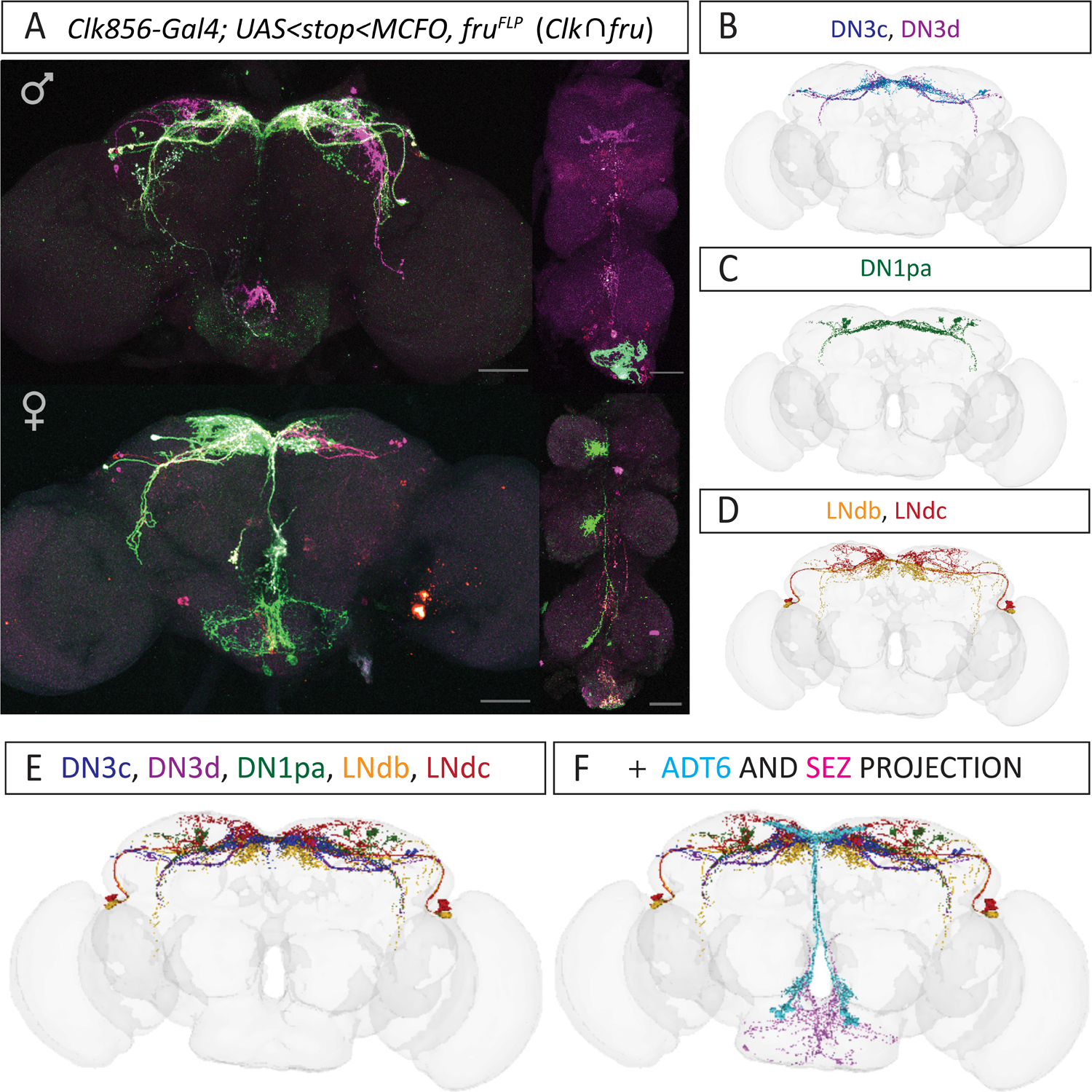
FlyWire connectome classification of *fru* ∩ *Clk* brain neurons. (**A**) Brain and VNC confocal maximum projections from 5–10 day old adults. *fru* ∩ *Clk* neurons are identified using the MultiColor FlpOut (MCFO; *UAS<Stop<MCFO*) reporters. Scale bars = 50 μm. Images were acquired with a 20X objective. The genotype is *w, Clk856-Gal4/+; UAS<Stop<MCFO, fru^FLP^/+* (**B-D**) FlyWire 3D projections of s-CPDN_3_C (DN3c), s-CPDN_3_D (DN3d), DN_1p_A (DN1pa), LN_d_B, and LN_d_C (LNdb and LNdc) neurons. The nomenclature is from the published circadian connectome ^19^. (**E**) Combined FlyWire projections of DN3c, DN3d, DN1pa, LNdb, and LNdc neurons. (**F**) Addition of FlyWire ADT6 and SEZ projections with DN3c, DN3d, DN1pa, LNdb, and LNdc neurons.

### FlyWire connectome classification of *fru* ∩ *Clk* brain neurons

Given our higher resolution morphological analyses of the *fru* ∩ *Clk* neurons with MCFO, we are able to classify them using the FlyWire connectome database, a circadian network connectome based on FlyWire, and annotation of *fru+* neurons in the FlyWire connectome ^18,19^. All the connectome neurons shown in Figure 1 are labeled as *fru+,* except the DN1pAs, providing further confidence in our assignments. The circadian connectome lists five types of DN3s. The *fru* ∩ *Clk* DN3s are likely DN3c (s-CPDN_3_C)^19^, and/or DN3d (s-CPDN_3_D)^19^, given they have projections in the protocerebral region, with some crossing the midline over the pars intercerebralis (PI). This is not observed for s-CPDN_3_B and s-CPDN_3_E neurons. We also rule out s-CPDN_3_A neurons because these have projections into the optic lobe, which we do not observe. For DN1s, we found the DN1pA neurons were most similar to *fru* ∩ *Clk* DN1 neurons. There was another classified set of DN1 neurons that have projections above the DN1 region that we did not observe in our MCFO staining. We propose the *fru* ∩ *Clk* LNds are LNdb and LNdc, and not the third category called LNda. The LNda have a large projection into the optic lobe that we did not observe. Using the FlyWire resource, we generated a composite of the neuronal classes (DN3c, DN3d, DN1pa, LNdb, and LNdc) that is similar to the *fru* ∩ *Clk* MCFO pattern (**Figure 1E**).

While neurons in the SEZ are not part of the defined ∼150 clock network, we identify them in the *fru* ∩ *Clk* MCFO staining. We observe median bundle neurons that project from the SEZ to the PI region that are similar in morphology to ADT6 neurons *(fru^+^)*, though we note that there over 230 ADT6 neurons that are difficult to distinguish by morphology. The cb1008 ADT6 neurons have similar cell numbers and projection. The cb1008 neurons also had projections near the DN3 and DN1 neurons, as we see in *fru* ∩ *Clk* MCFO staining, which gave us higher confidence. The projections we see in the SEZ were variable, however the overall cell body positioning and ring shape of the neurites is similar to an unlabeled neuron notated as cb0456 in FlyWire. Using the FlyWire resource, we generated a composite of all the potential neuronal classes that is highly similar to the *fru* ∩ *Clk* MCFO pattern (**Figure 1F**).

### Impact of the sex hierarchy on *fru* ∩ *Clk* neuron number

The *fru* ∩ *Clk* MCFO staining revealed that males had more DN3 neurons, compared to females. Scoring the number of *fru* ∩ *Clk* brain DN3 neurons blinded, showed that males have ∼6.75 and females have ∼3.75 (median counts; **Figure 2**). To determine if these differences are regulated by *fru*, we examined the number of *fru* ∩ *Clk* DN3 neurons when we overexpress different Fru^M^ isoforms that differ in their DNA binding domain (Fru^MA^, Fru^MB^, or Fru^MC^). Overexpression of Fru^MB^ lead to an increase in *fru* ∩ *Clk* DN3 female neurons compared to no overexpression, though genotype-matched males still had significantly more. Overexpression of Fru^MA^ lead to a decrease in *fru* ∩ *Clk* DN3 male neurons, resulting in no significant difference in neuron number between the sexes in that comparison. This result is consistent with known differences in functions of the Fru^M^ isoforms ^20–22^. When we examine a strong hypomorphic, transheterozygous *fru* allele combination that substantially reduces the amount of Fru^M^ protein (*fru^4-40^/fru^FLP^*; **Figure 2**)^23^, we find the number of *fru* ∩ *Clk* DN3 male neurons is reduced to the same number as female. We do not see this same result when we feminized the neurons by expressing Tra^F^, which is predicted to remove Fru^M^ from all neurons that express *Clk-Gal4*. The difference in the results may be because *fru^4-40^/fru^FLP^* results in a loss of Fru^M^ at any time Fru^M^ is produced, whereas the UAS-transgene manipulations are under the control of *Clk* expression, which may be at a later developmental time. Additionally, there could be a cell-nonautonomous requirement for *fru* for male *fru* ∩ *Clk* DN3 neurons. Across all these manipulations, we did not see an impact on neuronal morphology, consistent with the observation that we did not see gross morphological sexual dimorphism in *fru* ∩ *Clk* MCFO neurons.

**Figure 2.**
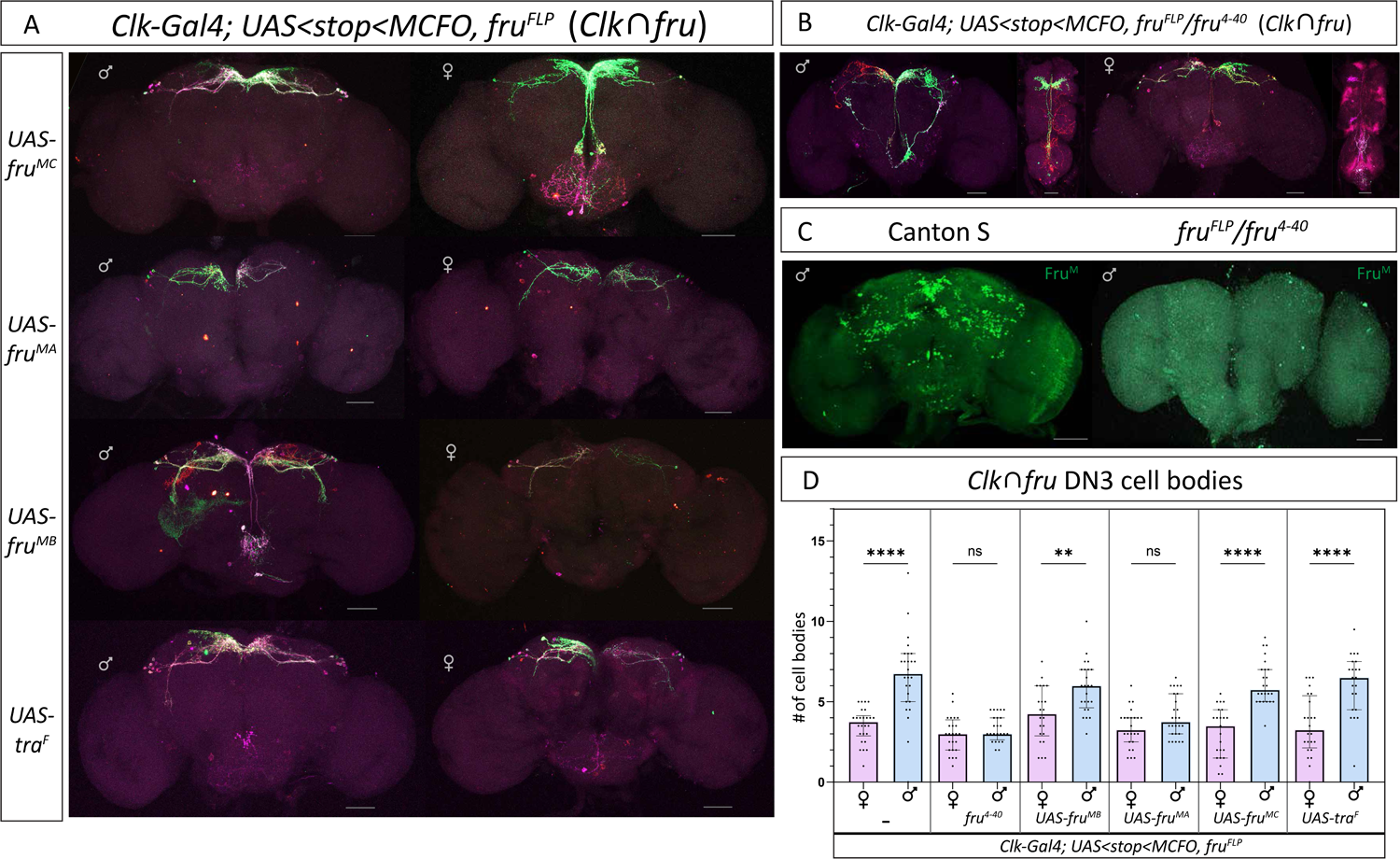
Impact of the sex hierarchy on *fru* ∩ *Clk* neuron number. (**A-B**) Brain and VNC confocal maximum projections from 5–10 day old adults. The genotype is: *w, Clk856-Gal4/+; UAS<Stop<MCFO, fru^FLP^/+*. Images show *fru* ∩ *Clk* neurons identified using the MultiColor FlpOut (MCFO) reporters. (**A**) MCFO straining with overexpression of Fru^M^ isoforms (Fru^MA^, Fru^MB^, or Fru^MC^) or Tra^F^ under UAS regulation. The genotype includes one of the UAS transgenes on the second chromosome. (**B**) MCFO staining in transheterozygous *fru* allele combination *fru^FLP^ / fru^4-40^*. (**C**) Brain confocal maximum projections from 5-10 day old adults. Wild type (Canton S strain) and *fru^FLP^* / *fru^4-40^* brains were stained with anti-Fru^M^ antibody. (**D**) Count of DN3 neurons in males and females at the ZT4-6 time window. Bar graphs show median with interquartile range. Two-tailed Mann-Whitney test comparing males and females for each effector. * = p <0.05, **=p<0.01, ***=p<0.001, **** = p<0.0001. *n*=22-26 hemibrains for each genotype examined, for both males and females. Scale bars = 50 μm. Images were acquired with a 20X objective.

### *fru* post-synaptic targets of circadian network neurons are dimorphic

To determine if there is additional sexual dimorphism in the circadian neural network, we identified the *fru* synaptic targets of *Clk-Gal4* neurons, using the trans-TANGO reporter system (**Figure 3**) ^15^. First, we restrict the trans-TANGO post-synaptic reporter to *fru* neurons, by using a reporter that relies on expression of *fru^FLP^* (green, **Figure 3A**). We observe sex-differences in the number and position of the *fru* synaptic targets, with more neurons in the male brain (**Figure 3C**). These *fru* synaptic targets are neurons in the posterior region of the brain with cell bodies near where the mushroom body cell bodies reside (here called midbrain cloud), ascending median bundle, and arch and lateral junction of the lateral protocerebral complex. If we examine individual brain regions, we find males more often have the midbrain cloud and the arch neurons detected (**Supplemental Figure 2**). Whereas in females we more often detect the ascending median bundle neurons (**Supplemental Figure 2**). We also detect more neurons in the ventral nerve cord in males (**Figure 3C**), in the VNC abdominal ganglion and VNC T2 midline neurons (**Figure 3A**). We also examined *fru* synaptic targets of *Clk-gal4* using a reporter for all *Clk-gal4* post-synaptic targets (**Figure 3B**; nuclear red) and detected their overlap with Fru^M^ (Figure 3B; nuclear purple Fru^M^ tagged with FLAG epitope, and green shows overlap). This approach revealed additional *fru* post-synaptic *Clk*-target neurons with cell bodies in the pars intercerebralis, though overall was consistent with the results using the trans-TANGO FLP-out reporter approach.

**Figure 3.**
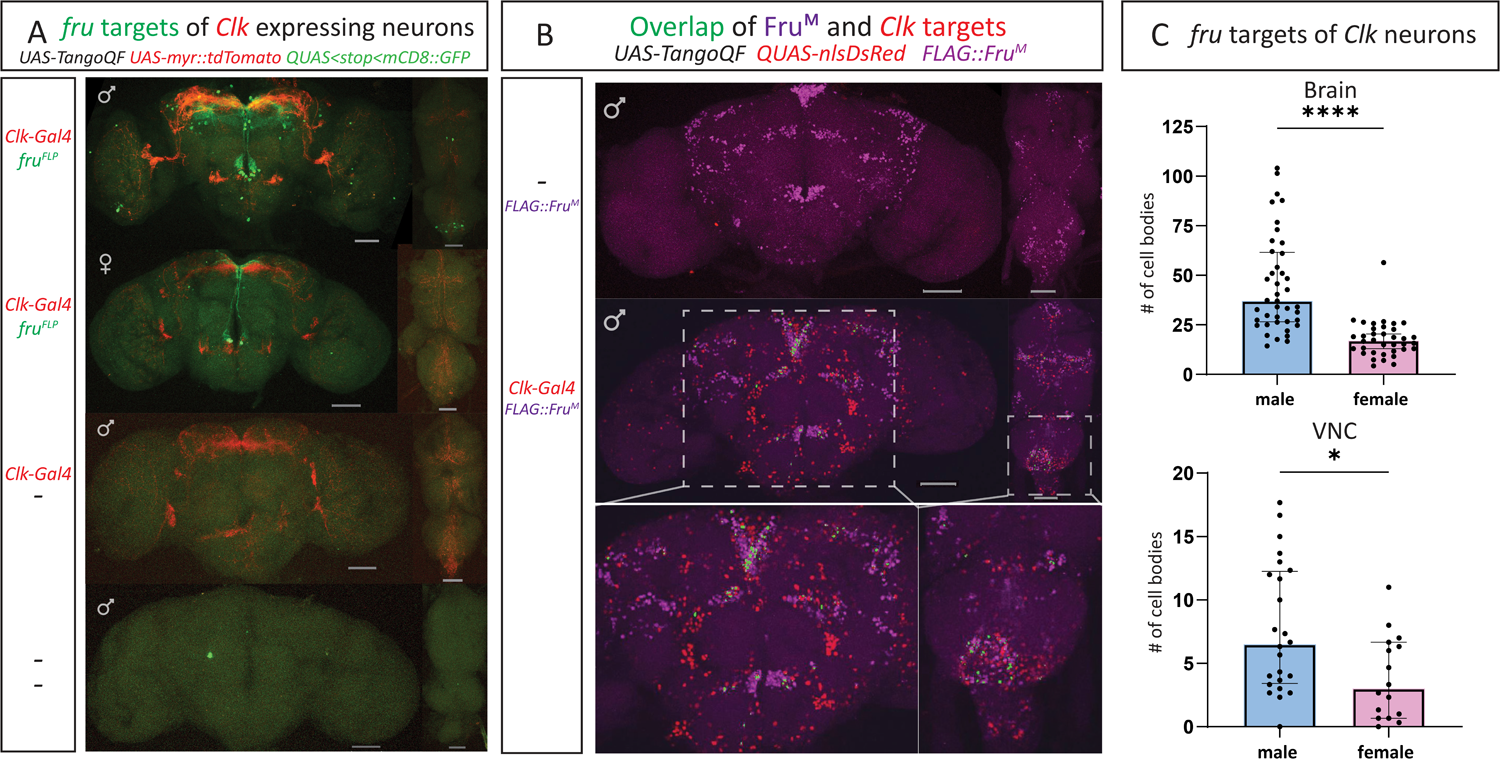
*fru* post-synaptic targets of circadian network neurons are dimorphic. (**A)** Brain and VNC confocal maximum projections from 5–10 day old adults. The trans-Tango reporter tool allows for visualization of *Clk856-Gal4* neurons (red), and *fru* synaptic targets of *Clk* (green). The genotype contains these elements: *trans-Tango/Clk856-Gal4; 10XUAS-IVS-myr::tdTomato, QUAS<stop<mCD8-GFP/ fru^FLP^*. The QF transcription factor is produced in the post-synaptic targets of *Clk*, with the QF-dependent reporter, *QUAS,* only active in *fru* neurons. Scale Bars = 50 μm. (**B**) Brain and VNC confocal maximum projections from 5–10 day old adults. The trans-Tango reporter tool allows for visualization of targets of *Clk856-Gal4* neurons (nuclear DsRed). These flies harbor a Crispr-modified *fru* allele that generates an amino-terminal Flag-tagged Fru^M^ (Flag::Fru^M^; anti-Flag purple). The genotype contains these elements: *5XQUAS-nlsDsRed;trans-Tango/Clk856-Gal4;Flag::Fru^M^*. Colocalization of the red and purple signal is shown in green. Scale Bars = 50 μm. (**C**) Male and female cell body counts of the post-synaptic *fru* targets of *Clk* neurons from (A), per brain and VNC. Bar graphs show median with interquartile range (IQR) and two-tailed Mann-Whitney test. Male Brain: *n*= 41 brains, IQR = 25-57, Range = 13-115; Female Brain: *n* = 36, IQR = 11-19, Range = 3-55, Male VNC: *n* = 24, IQR = 3.25-12, Range = 0-17; Female VNC: *n* = 16, IQR = 1-6.75, Range = 0-11. * = p <0.05, **** = p<0.0001

### *fru* ∩ *Clk* neuronal activity impacts circadian period and overall activity

Next, we determine the function of *fru* ∩ *Clk* neurons in circadian period length, by either activating (*UAS<stop<TrpA1;* hereafter called *TrpA1*) or silencing (*UAS<stop<Kir2.1;* hereafter called *Kir2.1*) *fru* ∩ *Clk* neurons (**Figure 4**), in strain background matched conditions. Similar to our previous study, in constant dark conditions, we found activating *fru* ∩ *Clk* neurons resulted in males having a shortened period length (23.2 hours), but females were similar to controls (23.8 hours for females; 23.8 hours for male and female controls). We were surprised that silencing *fru* ∩ *Clk* neurons also reduced circadian period in males (22.6 hours), and females (23.4 hours), compared to controls (23.8 hours for male and female controls) (**Figure 4B and Supplemental Table 1**). Males and females with either activated (*TrpA1*) or silenced (*Kir2.1*) *fru* ∩ *Clk* neurons had more overall activity compared to same-sex, strain-matched controls (*w^+^*), which could be related to the shortened circadian period. Females with activated *fru* ∩ *Clk* neurons (*TrpA1*) had the lowest circadian strength and percentage of rhythmic individuals (**Figure 4C and Supplemental Table 1**). The observation that continuous activation or silencing of *fru* ∩ *Clk* neurons had the same shortened circadian period phenotype in males, is consistent with a need for *fru* ∩ *Clk* neurons to change their activity across the day and night to maintain wild type period.

**Figure 4.**
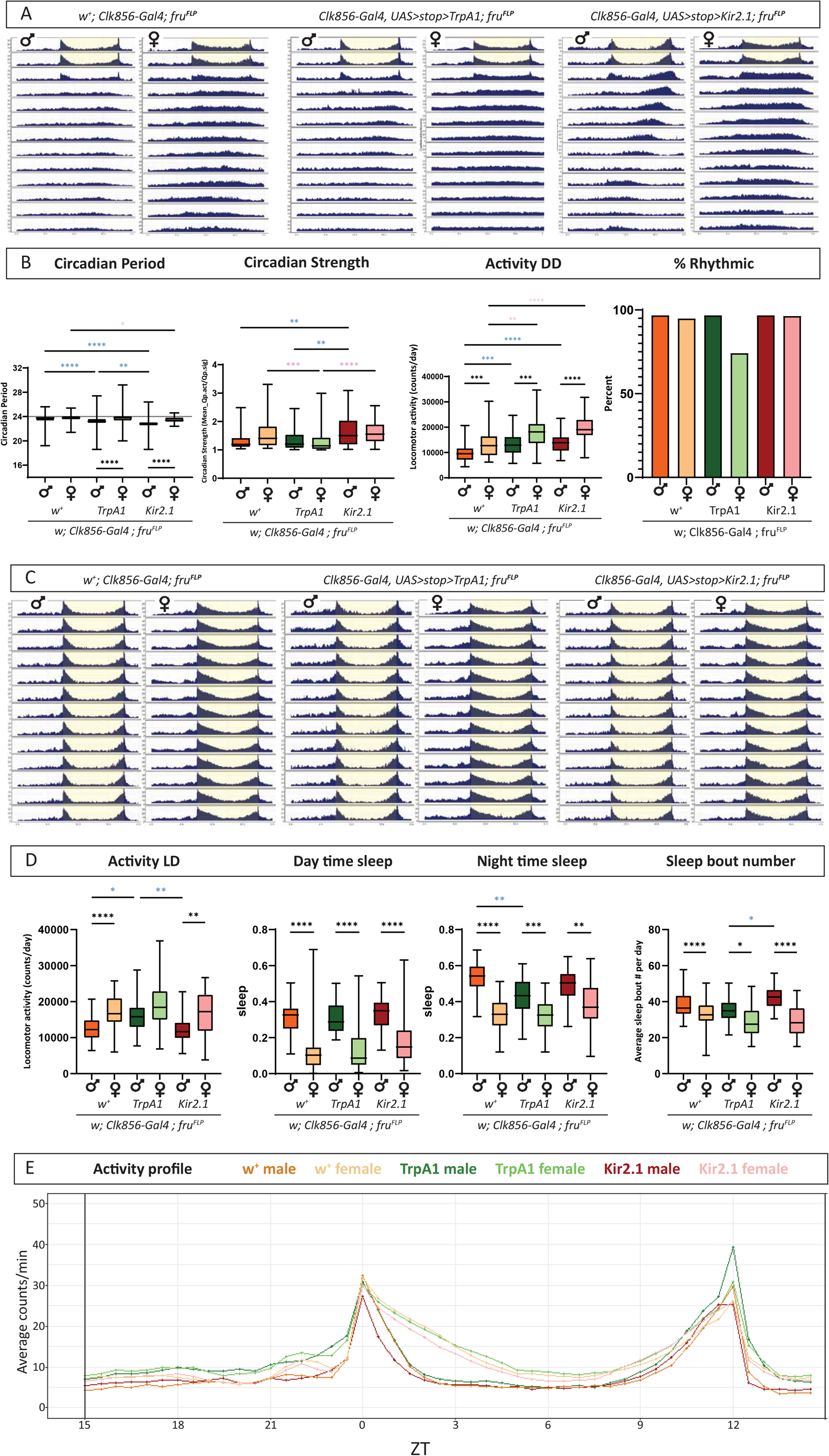
*fru* ∩ *Clk* neuron activity impacts circadian period and overall activity. **(A)** Actograms showing averaged activity over two days in light dark and eleven days in constant dark. The yellow shaded region indicates when lights are on at the end of light:dark entrainment (first two days in monitors). Males and females are 1) *w^+^, Clk856-Gal4/+; fru^FLP^* (controls, male n=62, female n=59), 2) *w; Clk856-Gal4/UAS<stop<TrpA1; fru^FLP^/+.* (TrpA1 activated, male *n*=62 female *n*=62), and 3) *w; Clk856-Gal4/ UAS<stop<Kir2.1; fru^FLP^/+.* (Kir2.1 silenced, male *n*=62 female *n*=55). Y axis range 0-35 counts. x axis. The data are plotted in 5-minute bins. (**B**) Circadian period, circadian strength (reported as Qp.act/Qp.sig.), average activity per day and number of rhythmic flies were calculated using ShinyR-DAM ^27^. % rhythmic reports ratio of flies with >1 Qp.act/Qp.sig. We performed a route method to find outliers, with a cut-off of Q of .01. Three flies were removed in total that had activity above 60,000 counts. **(C)** Actograms showing averaged activity over 13 days in LD, with the yellow shaded region indicating when lights are on. Males and females are 1) *w^+^, Clk856-Gal4/+; fru^FLP^* (male n=41, female n=42) 2) *w; Clk856-Gal4/UAS<stop<TrpA1; fru^FLP^/+.* (male *n*=38, female *n*=41), and 3) *w; Clk856-Gal4/ UAS<stop<Kir2.1; fru^FLP^/+.* (male *n*=42 female *n*=38). **(D)** Activity across 13 days of LD activity, sleep between ZT0 and ZT12 (lights on) indicated as daytime sleep, and sleep between ZT12-ZT0 (lights off) indicated as nighttime sleep, and average # of sleep bouts/day averaged across all 13 days of LD activity. In **B** and **D** the box-whiskers show min-max tails. Kruskal-Wallis with Dunn’s multiple comparisons for statistical analysis. Black asterisks indicate sex-differences differences within genotype, blue asterisks indicate differences across male genotypes, and pink asterisks indicate differences across female genotypes. Tests between males and females of different genotypes were not performed. * = p <0.05, **=p<0.01, ***=p<0.001, **** = p<0.0001. (E) Averaged activity profile of conditions when *fru* ∩ *Clk* neurons are activated (TrpA1), or silenced (Kir2.1) from 13 days of LD averaged into 30-minute bins.

We also examined activity and sleep when *fru* ∩ *Clk* neurons are either activated (*TrpA1*) or silenced (*Kir2.1*), in 12 hours light and 12 hours dark conditions (LD), using strain background matched conditions. Control females are more active than males (*w^+^*) in LD (**Figure 4D**). When *fru* ∩ *Clk* neurons are activated (*TrpA1*) this sex difference is no longer observed, with males having increased LD activity. This may be due to the reduction of nighttime sleep in males when *fru* ∩ *Clk* neurons are activated (*TrpA1*), as compared to controls (*w^+^*). Across all three conditions, males sleep more than females, during the day and night. Activating *fru* ∩ *Clk* neurons resulted in fewer sleep bouts, as compared to silencing *fru* ∩ *Clk* neurons, which is a potential explanation for the reduced sleep at night when *fru* ∩ *Clk* neurons are activated (**Figure 4D**). To determine if there are structural differences in activity in LD, we examined averaged activity data across day and night.

Males of all three genotypes show a rapid reduction in activity after lights on (ZT 0), compared to females, and a longer afternoon siesta, as previously reported ^1^. One trend in the data is that male (red; **Figure 4E**) and females (pink; **Figure 4E**) with *fru* ∩ *Clk* neurons silenced *(Kir2.1)* reduced their activity more quickly after lights on. Consistent with our observation that males with activated *fru* ∩ *Clk* neurons (*TrpA1*) having reduced nighttime sleep, these males show a much larger peak in activity after lights off (dark green; ZT12; **Figure 4E**) and continue to remain more active during the night.

### Impact of *fru* ∩ *Clk* neuron number and sex-specific identity on circadian period

Given the *fru^4-40^/fru^FLP^* genetic background resulted in a loss of sexual dimorphism in *fru* ∩ *Clk* DN3 neuron number (**Figure 2**), we could determine if the sex-difference in DN3 neuron number is responsible for the reduction in circadian period length in males with activated or silenced *fru* ∩ *Clk* neurons (**Figure 4**). To better understand the *fru^4-40^/fru^FLP^* transheterozygous allele combination, we tested males in courtship assays. Across all metrics *fru^4-40^/fru^FLP^* transheterozygotes had reduced courtship activity as compared to wild type males, though they still courted (**Figure 5**). This indicates that the *fru^FLP^* allele is a partial loss of function, despite the lack of Fru^M^ staining we observed in *fru^4-40^/fru^FLP^* (**Figure 2**), as previously suggested ^24^. Unexpectedly, the *fru^4-40^/fru^FLP^* males and females both show a reduced circadian period when *fru* ∩ *Clk* neurons are activated or silenced, indicating that the large *fru^4-40^* deficiency has a non-sex-specific genetic interaction causing a reduced circadian period (**Figure 5**). Therefore, it is not clear if the reasons males show a reduced circadian period when *fru* ∩ *Clk* neurons are either activated or silenced is due to differences in neuron number.

**Figure 5.**
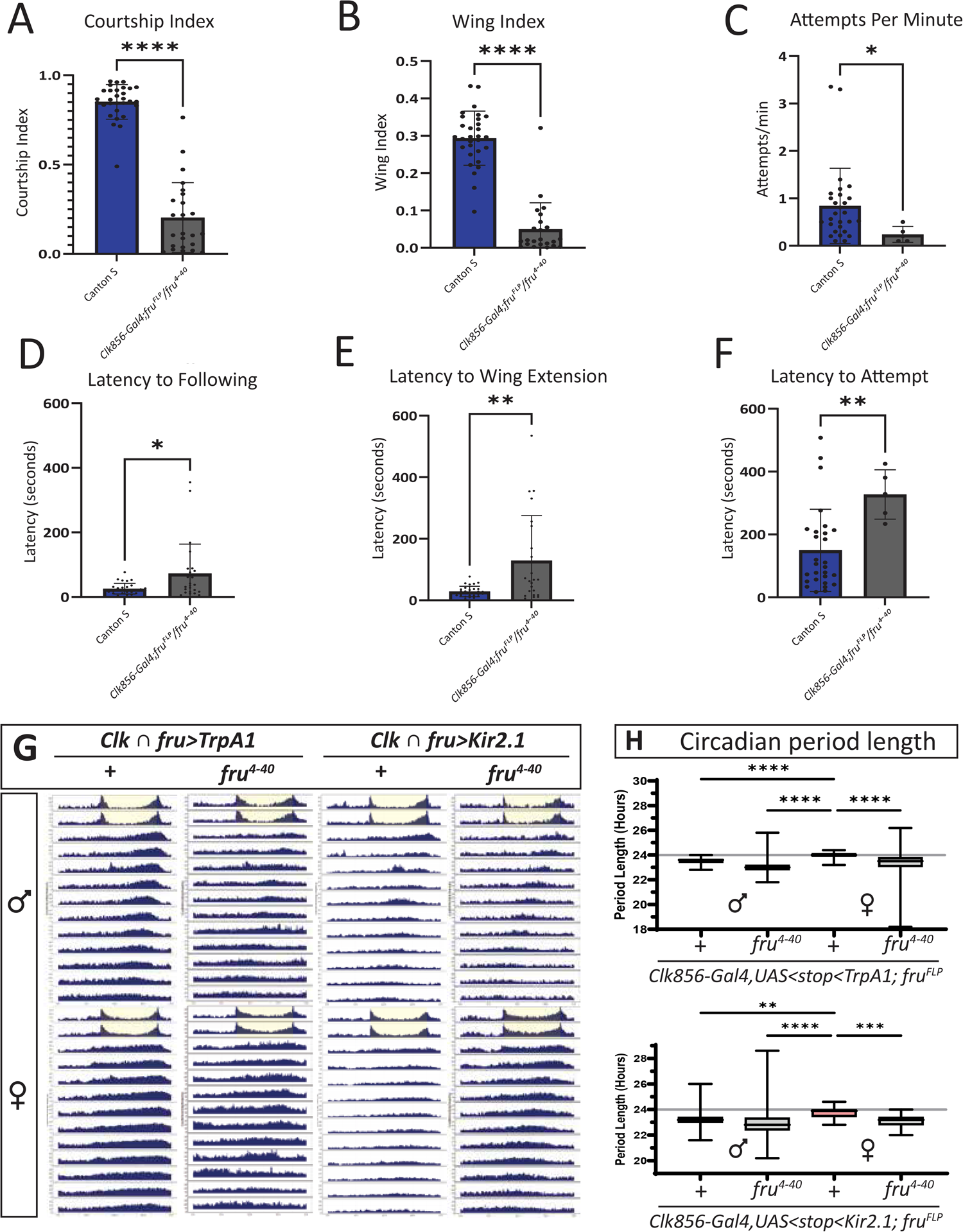
Impact of *fru* ∩ *Clk* neuron number and sex-specific identity on circadian period. (**A-F**) Male courtship analyses of Clk856-Gal4/+; *fru^FLP^ / fru^4-40^* and wild type Canton S. Courtship data shows mean ± standard deviation, with Mann-Whitney statistical tests. *n* = 30 per genotype (**A**) Courtship index measured as the time spent following divided by the total observation time. (**B**) Wing index measured as the time spent wing extending divided by the total observation time. (**C**) Number of attempts per minute of observation time. (**D**) Latency to following. (**E**) Latency to wing extension. (**F**) Latency to attempt. (**G**) Actograms showing averaged activity over two days in light:dark (first two days in monitors) and eleven days in constant dark. The yellow shaded region indicates when lights are on at the end of light:dark entrainment (first two days of actogram). Y-axis range 0-35 counts. X-axis is plotted in 5-minute bins. (**H**) Circadian period calculated from ShinyR-DAM. Box and whiskers plots with min to max of the data. Males and females of *Clk856-Gal4/UAS<stop<TrpA1; fru^FLP^*. Male *n*=28, female *n* =30. *Clk856-Gal4/UAS<stop<TrpA1; fru^FLP^/fru^4-40^*. Male *n* =23 female *n* =28. *Clk856-Gal4/UAS<stop<Kir2.1; fru^FLP^*. Male *n* =25, female *n* =28. *Clk856-Gal4/UAS<stop<Kir2.1; fru^FLP^/fru^4-40^*. Male *n* =24, female *n* =31. Kruskal-Wallis with Dunn’s multiple comparisons for statistical analysis. For all analyses, * = p <0.05, **=p<0.01, ***=p<0.001, **** = p<0.0001.

To further determine if sex-specific identity impacted the phenotype we observed when *fru* ∩ *Clk* neurons are either activated or silenced we introduced transgenes that either feminize (*UAS-Tra^F^*) or masculinize (*UAS-tra* and *UAS-tra-2* RNAi) neurons and did not see an impact on the phenotype (**Supplemental Table 1**). In addition, we show that the male-specific reduced circadian phenotype is due to *fru* ∩ *Clk* neurons in the brain, by introducing a transgene that blocks *Clk-Gal4* activity in the VNC (*tsh-Gal80*) (**Supplemental Figure 3**) ^25^.

### *fru* ∩ *Clk* neuronal activity has impacts on reproductive behavior

To determine the role of *fru* ∩ *Clk* neurons in courtship, we examined courtship behavior when these neurons are either activated or silenced. While the TrpA1 channel is temperature gated, all the circadian phenotypes we observed were in 25° C conditions. It is also known that the TrpA1 channel has temperature-independent activation ^26^. For consistent comparisons, we test for courtship changes in the same conditions we saw circadian phenotypes. We saw a significant increase in overall courtship when the neurons are activated (courtship index; *TrpA1*), and an increase in the number of copulation attempts/per minute, suggesting that *fru* ∩ *Clk* neurons have a role in courtship timing or gating.

To determine if mating status would reveal additional roles for *fru* ∩ *Clk* neurons, we examined courtship, mating, activity and sleep in males and females after mating. Post-mating, males and females did not have differences in courtship and remating metrics, time to next mating, sleep or overall activity when *fru* ∩ *Clk* neurons are either activated or silenced (**Supplemental Figure 4**). Across all genotypes, mated females show a decrease in sleep, compared to unmated controls. Activating or silencing the *fru* ∩ *Clk* neurons did not reveal post-mating differences in sleep compared to control *(w^+^)*, though mated females with silenced *fru* ∩ *Clk* neurons *(Kir2.1)* had less overall activity compared to mated controls *(w^+^)*. Therefore, *fru* ∩ *Clk* neurons do not appear to be important for post-mating behavioral changes.

### Period cycling is shifted due to changes in *fru* ∩ *Clk* neuron activity

Next, to understand how the shortened circadian period arises when *fru* ∩ *Clk* neuron activity is altered, we examined Per cycling in the critical sLNvs clock pacemaker neurons. Per protein immunostaining across 6 timepoints, starting seven days after the initiation of constant dark conditions reveals a similar phase shift when *fru* ∩ *Clk* neurons are either activated or silenced, consistent with the change in behavior (**Figure 7**). We expect to observe a ∼7-8 hour shift in Per cycling after 7-8 days in DD. The *fru* ∩ *Clk* neurons (DN1, DN3, and LNds) are thought to be downstream of the sLNv pacemaker neurons. This suggests that there is feedback to the sLNvs either through synaptic connections, or indirectly, via the changes in behavior that are directed by *fru* ∩ *Clk* neurons. It has been shown that sLNvs are a target of DN1, and that DN1 has robust sex differences in activity ^5^. Our results provide further evidence that DN1s may feedback on the core pacemaker sLNvs to change rhythmic behaviors.

## Discussion

We show that the *fru* ∩ *Clk* neurons direct sex-specific changes in circadian period length and impact male courtship behavior. That both activating or silencing the *fru* ∩ *Clk* neurons results in a shorter period length in males, suggests that activity modulation in these neurons is important for maintenance of the circadian period. We found that males have more *fru* ∩ *Clk* DN3 neurons, which may account for the stronger male circadian phenotype. An examination of the *fru* ∩ *Clk* neurons from our annotation based on the connectome, and a more focused analyses on the functions of *fru* ∩ *Clk* subsets will help address the basis for the sex-difference. While a lot is known about the clock, this study provides additional annotation of the DN3 clock neurons, which is a class that is not well understood. Why the circadian period length in males is more sensitive to changes in *fru* ∩ *Clk* neuronal activity and which neurons direct this difference is an interesting next question. Additionally, it is important to understand if there are additional behavioral outputs these neurons could impact in different behavioral paradigms. Our labeling of *fru^+^* post-synaptic targets of the clock also show additional sexual dimorphism downstream of the clock that is important to understand.

We found a link between the clock and courtship, with activation of *fru* ∩ *Clk* neurons changing the rate of copulation attempts. This suggests a role for *fru* ∩ *Clk* neurons in courtship gating or timing. Future studies to understand if the molecular clock resides in the *fru* circuit to direct additional rhythmic courtship behaviors is an important future goal. For example, examining another rhythmic courtship behavior, such as wing song, may yield additional roles for *fru* ∩ *Clk* neurons. Overall, this study provides an additional model to understand how neural networks that have a role in distinct behaviors can cross-influence behavioral outcomes with co-expression of the master regulatory transcription factors an important step.

## Acknowledgments

We thank the Bloomington Drosophila Stock Center (NIH P40OD018537) and Drosophila colleagues for generously providing stocks and antibodies. Several antibodies used in this study were obtained from the Developmental Studies Hybridoma Bank, created by the NICHD of the NIH and maintained at The University of Iowa, Department of Biology, Iowa City, IA 52242. We used FlyBase to find information about genes, stocks, phenotypes and function^28^. We also thank Colleen Palmateer for R code and helpful input. This work was supported by NIH MIRA grant 5R35GM145282 and NIH R01GM073039 and R01GM116998.

## Author contributions

Investigation, A.D., B.B., D.K.P., M.A.K., M.T., and M.N.A.; Formal Analysis, A.D., B.B., D.K.P., M.A.K., M.T., and M.N.A; Conceptualization, M.N.A.; Writing-Original Draft, A.D., B.B., and M.A.K.; Writing – Review & Editing, A.D., B.B., M.A.K., M.T., and M.N.A.; Visualization. A.D., B.B., D.K.P., and M.A.K.; Supervision, M.N.A.; Project Administration, M.N.A.; Funding Acquisition, M.N.A.

## Declaration of interests

The authors declare no competing interests.

## Material Availability Statement

All unique/stable reagents generated in this study are available from the lead contact without restriction.

## Data and code availability

Source data for all figures is available from the lead contact without restriction.

## Supplemental information titles and legends

**Document S1. Figures S1-S5**

**Supplemental Figure 1. *fru* ∩ *Clk* neuronal projections at different time points in a 24 hour period in light:dark conditions.**

**Supplemental Figure 2. Trans-Tango brain regions and quantification**

**Supplemental Figure 3: Validation of tra RNAi, tra2 RNAi and tsh-gal80 transgenes**

**Supplemental Figure 4: Sleep and activity post-mating**

**Supplemental Figure 5. Per time course images**

**Supplemental Table 1: Excel file containing Circadian Period Length and % rhythmic.**

## Methods

### Drosophila husbandry and genetic strains

All strains, unless otherwise indicated, are aged in humidified incubators at 25°C on a 12-hr light and 12-hr dark cycle. The laboratory Drosophila media composition is: 33 l H2O, 237 g agar, 825 g dried deactivated yeast, 1560 g cornmeal, 3300 g dextrose, 52.5 g Tegosept in 270 ml 95% ethanol and 60 ml propionic acid. To control for impacts of strain background on several of the behavioral studies the following genes/transgenes were introgressed for five generations into a common *w^1118^* genetic background: *w^+^* (control), *UAS<stop<TraA1::myc*, and *UAS<stop<Kir2.1*. Virgin females from these strains were crossed to *Clk856-Gal4*; *fru^FLP^* males to perform experiments on circadian period length (Figure 4), activity and sleep (Figure 4), and courtship (Figure 6). All flies in these experiments had wild type eye color.

**Figure 6.**
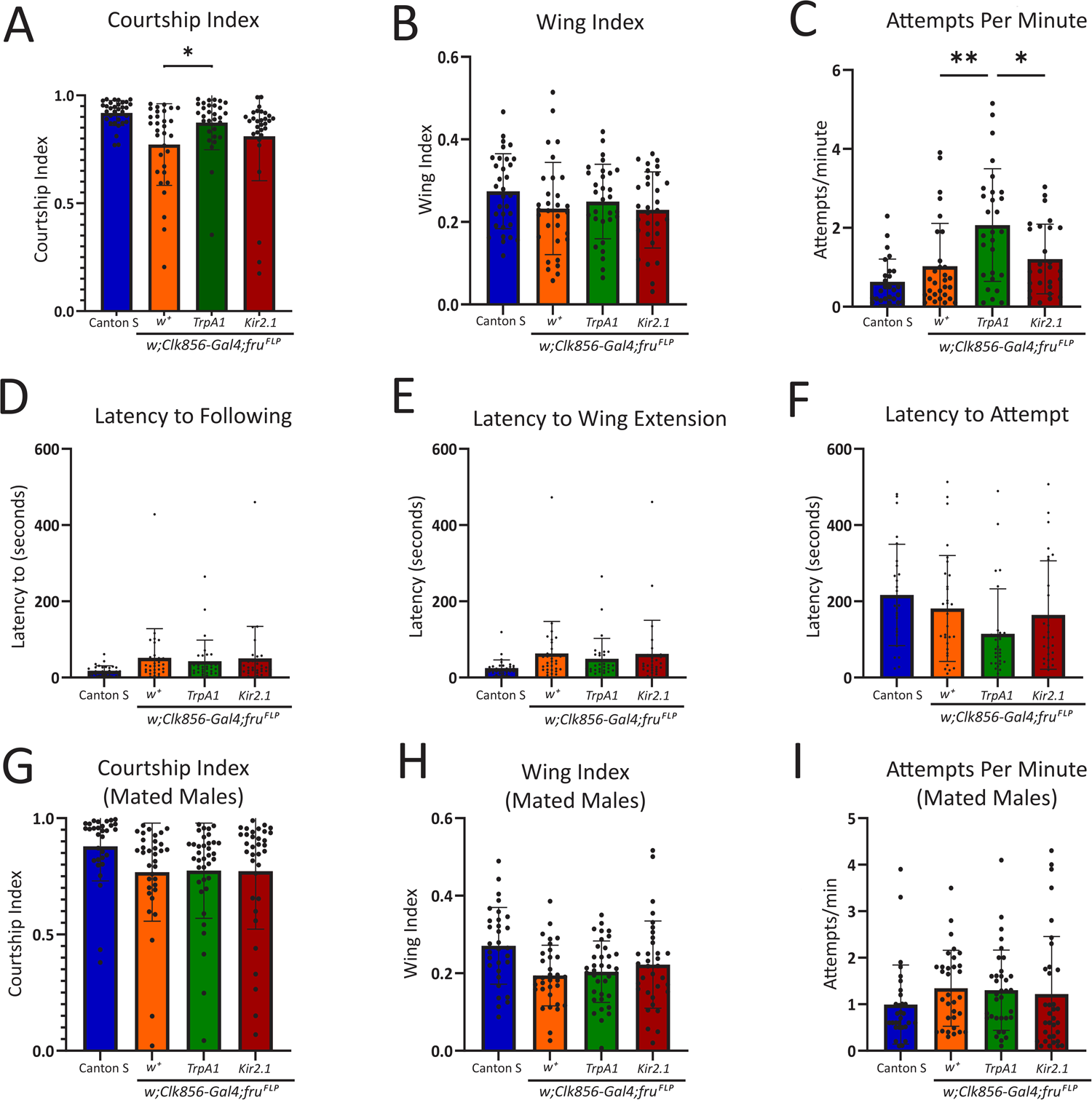
*fru* ∩ *Clk* neuronal activity has impacts on reproductive behavior. **(A-I)** Analysis of male courtship when *fru* ∩ *Clk* neuronal activity is altered. Courtship data shows mean ± standard deviation, with Kruskal-Wallis or one-way ANOVA statistical tests with multiple comparisons. *n*= 32 per genotype**. (A)** Courtship index measured as the time spent following divided by the total observation time. Kruskal Wallis with Dunn’s multiple comparisons for statistical analysis. **(B)** Wing index measured as the time spent wing extending divided by the total observation time. One-way ANOVA with Tukey’s multiple comparisons for statistical test. **(C)** Number of attempts per minute of observation time. One-way ANOVA with Tukey’s multiple comparisons for statistical test. **(D)** Latency to following in seconds. **(E)** Latency to wing extension in seconds. Kruskal Wallis with Dunn’s multiple comparisons for statistical analysis. **(F)** Latency to attempt in seconds. Kruskal Wallis with Dunn’s multiple comparisons for statistical analysis. **(G-I)** Courtship indices of males that were placed in the courtship assay following an initial mating. *n*= 31 CS, 35 *w^+^,* 36 *TrpA1* and 35 *Kir2.1.* Kruskal Wallis with Dunn’s multiple comparisons for statistical analysis. **(G)** Courtship index **(H)** Wing extension index. Kruskal Wallis with Dunn’s multiple comparisons for statistical analysis. **(I)** Attempts per minute. Kruskal Wallis with Dunn’s multiple comparisons for statistical analysis. For all analyses, * = p <0.05, **=p<0.01, ***=p<0.001, **** = p<0.0001.

**Figure 7:**
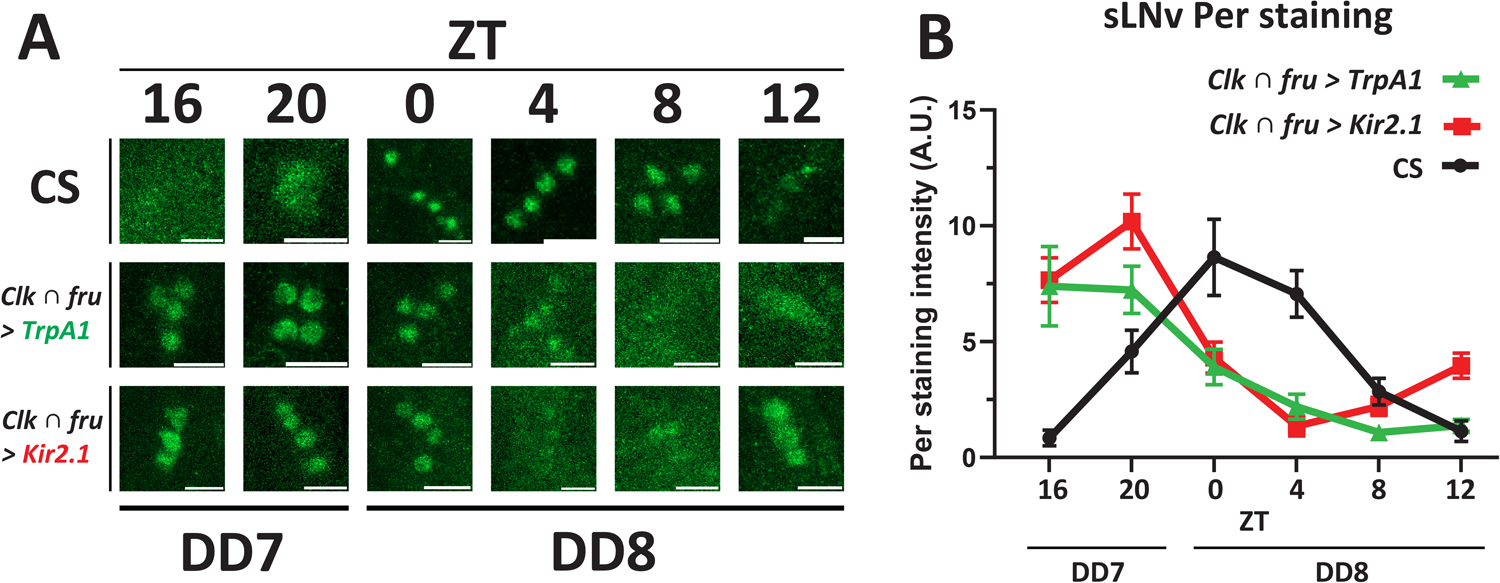
Behavioral rhythms in constant dark are reflected in sLNv Per protein cycling. (**A**) Representative images of anti-Per staining in sLNv neurons across 6 time points. Scale bars indicate 10um. (**B**) Quantification of Per staining intensity. Values = [(Nuclear Signal – Background) / Background]. Mean ±95% CI. *n* = 14-70 cells, (5-22 hemibrains, 4-11 brains). Experimental genotypes are: Canton S, *w*; *Clk856-Gal4, UAS<Stop<TrpA1/+; fru^FLP^/+* and *w; Clk856-Gal4; UAS<Stop<Kir2.1/+; fru^FLP^/+*

### Immunostaining and microscopy

Adult brains and VNCs were dissected and imaged as previously described^3^. Both primary and secondary antibodies were diluted in TNT (Tris-NaCL-Triton buffer; 0.1 M Tris–HCl, 0.3 M NaCl, 0.5% Triton X-100). All confocal microscopy was performed on a Zeiss LSM 900 system, with Zeiss Plan-Apochromat 20x/0.8 and 63x/1.4 objectives. The z-stack slice interval for all images was 1.0μm or 2.0μm. A 1 Airy Unit (AU) pinhole size was selected in Zeiss software (ZEN Blue 3.6) for each laser line: 488 nm: 38 μm; 555 nm: 34 μm; and 639 nm: 39 μm. All images were acquired in 16-bit grayscale space at 1024 × 1024 or 512 x 512 pixel resolution (0.19x or 0.39x Nyquist Sampling) with bidirectional scanning. The primary antibodies are rabbit α-Per (1:500; gift from Michael Rosbash), rat α-Fru^M^ (1:200) ^29^, mouse α-flag (1:500; Sigma, F1804), rabbit α-HA-tag (1:300; Cell Signaling, 3724S), Chicken α-Myc (1:1000; Invitrogen, A21281), Mouse α-PDF (1:1000; DSHB, PDF C7). The secondary antibodies are: rabbit α-GFP 488 (for Tango with flp-out; 1:500; Invitrogen, A21311), goat α-rat 488 (for anti-Fru; 1:500; Invitrogen, A11006), rabbit α-V5 tag 549 (for MCFO,1:500; Rockland, 600-442-378), Goat α-rabbit 633 (for MCFO, 1:500; Invitrogen, A21071), Goat α-mouse 488 (for MCFO,1:500; Invitrogen, A11001), goat α-mouse 633 (for Flag, 1:500; Invitrogen, A21052), Goat α-rabbit 488 (for Per staining,1:500; Invitrogen, 32731), Goat α-chicken 546 (for myc, 1:500; Invitrogen, A11040), Goat α-mouse 633 (for Pdf, 1:500; Invitrogen, A21052)

### Quantification of neuron numbers

To quantify *fru* ∩ *Clk* MCFO neurons and the trans-Tango post-synaptic *fru* targets, confocal images were blinded, and cell bodies were counted independently by 2 or 3 people using Zeiss microscopy software ZEN Blue 3.6. Statistical tests were performed on averaged cell body counts across the independent observers. Results of statistical tests were consistent across averaged counts and within each observer’s counts. Number of samples used for cell-counting data is reported in the results sections.

### Circadian Assays

Circadian assays were performed using Trikinetics DAM5H activity monitor (Trikinetics, Waltham, MA) and all data was collected in 1 minute bins using DAM System 311. Flies were collected Monday – Wednesday and loaded into 5×65mm glass tubes that contained our standard laboratory food. Tubes containing the flies were kept in a LD 12:12 humidified incubator until being loaded into DAM5H on Friday. The incubator was sealed with tin foil to prevent any light leaking in. The light:dark (LD) assay was conducted at 25°C on a 12-hr:12-hr LD cycle and beam cross activity was recorded in one min bins for 15 days. Incubator lights turned on at 9 am and off at 9 pm. This resulted in 13 days of data analyzed for all genotypes in the LD assay. In the dark:dark (DD) assay, flies were first entrained and aged at 25°C on a 12-hr:12-hr LD cycle for 5-7 days, allowing for 2 days of LD data for analysis. Next, we switched to a 12-hr:12-hr DD cycle for 12 days allowing for 11 days of data analyzed for DD conditions and circadian period (constant darkness for 12 days). Beam cross activity for the DD assay was recorded in 1-min bins. Data were analyzed using ShinyR-DAM for both assays, excluding the first and last day the flies were placed in the incubator. Dead flies were considered those with less than 50 beam cross events per day and were removed from the analyses. ShinyR-DAM analysis of sleep was only performed on LD assay data. ShinyR-DAM measures sleep events using a 5-min sliding window, where 5 min of inactivity is considered a sleep event. Circadian period analysis was performed in ShinyR-DAM using the DD assay data. The default ShinyR-DAM parameters were used as follows: Chi-Sq testing range of 18–30 hr, a Chi-Sq period testing resolution of 0.2, and a rhythmicity (circadian strength) threshold for filtering arhythmic individuals (Qp.act/Qp.sig) of 1.

### Per time series

0-24 hour-old control and experimental flies were collected and individually housed on days 1-4. On day 3-4 they began entrainment in LD (12:12) at 25°C. After 6-9 days of entrainment, the flies were placed in DD for 7-8 days until dissection on day 7-8. Flies were 14-16 days old at dissection. For DD conditions, individually housed flies were wrapped in aluminum foil, placed in light-proof mylar bags, and kept in a dark incubator at 25°C for 7-8 days until dissection. Incubator LD entrainment times were varied while dissections were performed at the same time of day, resulting in different zeitgeber times at dissection. For each dissection, individual vials of 10-15 flies were removed from the light-proof bags, aluminum foil unwrapped, and flies were anesthetized on a CO_2_ pad. Flies were euthanized and permeabilized in 100% Ethanol for 2-10 seconds, washed in PBS, and fixed in 4% Paraformaldehyde in PBS (PFA) for 15 minutes. Fly heads were decapitated during PFA incubation. Brains were then dissected and fixed for an additional 10 minutes in 4% PFA and processed using our standard immunohistochemistry protocol, as described previously. Brains were incubated overnight at 4°C in primary antibodies, then washed in TNT and incubated in secondary antibody for 4-6 hours at room temperature (18-22°C) with gentle rocking. Brains were scanned on the confocal microscope with 20x air and 63x oil immersion objectives.

### Immunostaining quantification for Per time series

Confocal settings were kept consistent across all scans. Confocal scans were captured in 16-bit grayscale space using a 63x oil immersion lens at 0.45x digital zoom with bidirectional scanning and 1um or 2um z-slices. Pixel resolution was 512×512 or 1024×1024 (0.19x or 0.39x Nyquist Sampling). Confocal scan files (.czi) were quantified in ImageJ (Fiji) using the Bio-Formats Importer plugin. Individual cells of circadian neuron populations were identified based on location, morphology, and intensity of the PDF, PER, and myc signals. Nuclear regions of interest (ROIs) were drawn manually on a single slice of the z-stack for each cell of interest (1 cell = 1 ROI = 1 slice). Average Per staining pixel intensity was measured from nuclear ROIs. Average background pixel intensity was manually drawn as a representative region surrounding the cell population. Normalized Values = (Nuclear Signal – Background) / Background. Within hemibrains, each cell of a population was measured against a single common background for that population. Representative images were processed and exported using ImageJ. Maximum intensity projections were generated of z-stacks for circadian neuron populations. “Auto” Brightness and Contrast settings were applied to each channel of maximum projections in ImageJ. Scale bars = 10um.

### Courtship and mating assays

For male courtship assays, all male flies were collected within 0-6 hours post-eclosion and aged individually for 5-10 days in an incubator at 25°C on a 12-hr light/12-hr dark cycle. Female Canton S virgin flies were collected, and group housed in groups of ∼10 in an incubator at 25°C on a 12-hr light/ 12-hr dark cycle for 5-10 days. The assays were performed using a 10mm 8-well courtship chamber set on a 25°C -temperature block between ZT5 to ZT9. Courtship activity was recorded for 10 minutes or until successful copulation (n=∼30). Video recordings were analyzed blinded using Nodulus Observer XT software. Courtship Index and Wing Index were measured by dividing the time the experimental male fly spent doing each respective behavior toward the female by the total observation time. Attempted copulations per minute, successful copulations and latency to all courtship behaviors were also recorded per experimental male genotype. For male remating assays, males were placed in a mini-vial with one virgin Canton S at ZT5-8 and allowed to mate. Males that mated were immediately assayed with a new virgin in courtship chambers, as above.

For female mating assays all female flies were collected as virgins and aged in groups of ∼10 in an incubator at 25°C on a 12-hr light/12-hr dark cycle for 5-10 days. The Canton S males were individually aged as above for courtship. Latency to successful copulation was measured in courtship chambers, as described for male mating. For female remating assays, virgins of the experimental genotypes were single housed in mini-vials, as were Canton S males, and aged for 5-10 days 25°C on a 12-hr light/12-hr dark cycle. At ZT0-2 males were added to the female vials and mating was observed.

GraphPad Prism was used to conduct statistical analysis and statistical significance (p<0.05) using nonparametric Kruskal-Wallis ANOVA test with Dunn’s post-hoc correction for courtship index, all latency data and all remating data. A One-Way ANOVA with Tukey’s Multiple Comparisons was conducted on wing index and attempts per minute data from our initial mating assay. The *fru^4-40^* courtship data was analyzed in Prism using a Mann-Whitney U Test.

### Post-mating Locomotor Assay

Males 0-6 hours post-eclosion and virgin females were collected and aged individually for 5-10 days in mini-vials in an incubator at 25°C on a 12-hr light: 12-hr dark cycle. Matings were done in mini-vials from ZT0-ZT2 with the males or females of the experimental genotype being mated to a Canton S female or male, respectively. Following mating, both mated and unmated flies of the experimental genotype were placed individually in circadian vials, without anesthesia. Circadian vials were put into a DAM system to monitor activity immediately after mating, under LD conditions. Activity counts were collected starting at ZT 3 for 13 hours. For daytime sleep, activity of mated females was analyzed from ZT 3-12 on day 1. For nighttime sleep, activity of mated females was analyzed from ZT12-15 on day 1, using a 1-minute sliding window analysis in excel to find 5-minute windows of inactivity. ShinyR-DAM was used to analyze activity from ZT 3-15 on day 1.

### Key Resources Table

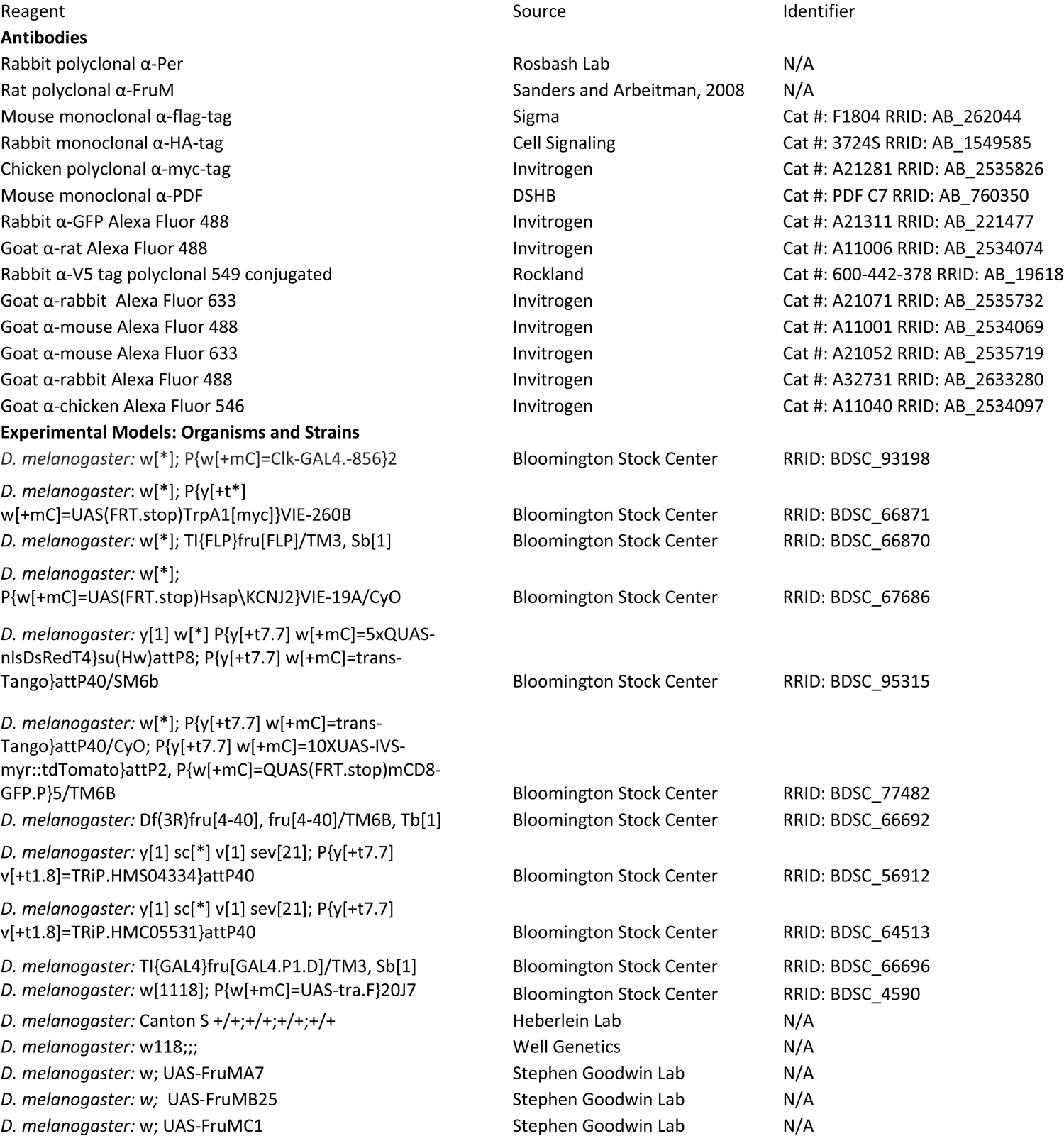
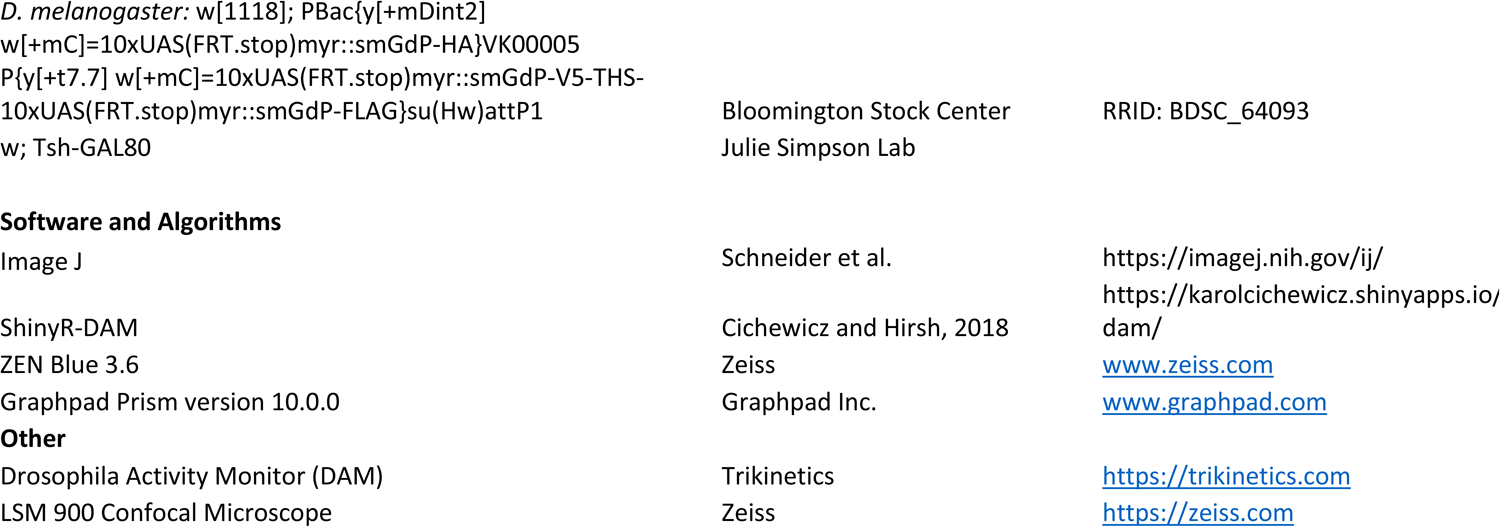

## Notes

### Competing Interest Statement

The authors have declared no competing interest.

## Bibliography

1. Dubowy, C., and Sehgal, A. (2017). Circadian Rhythms and Sleep in Drosophila melanogaster. Genetics 205, 1373–1397. 10.1534/genetics.115.185157.

2. Asahina, K. (2018). Sex differences in Drosophila behavior: Qualitative and Quantitative Dimorphism. Curr Opin Physiol 6, 35–45. 10.1016/j.cophys.2018.04.004.

3. Palmateer, C.M., Artikis, C., Brovero, S.G., Friedman, B., Gresham, A., and Arbeitman, M.N. (2023). Single-cell transcriptome profiles of -expressing neurons from both sexes. Elife 12. ARTN e7851110.7554/eLife.78511.

4. Cavanaugh, D.J., Vigderman, A.S., Dean, T., Garbe, D.S., and Sehgal, A. (2016). The Drosophila Circadian Clock Gates Sleep through Time-of-Day Dependent Modulation of Sleep-Promoting Neurons. Sleep 39, 345–356. 10.5665/sleep.5442.

5. Guo, F., Yu, J., Jung, H.J., Abruzzi, K.C., Luo, W., Griffith, L.C., and Rosbash, M. (2016). Circadian neuron feedback controls the Drosophila sleep--activity profile. Nature 536, 292–297. 10.1038/nature19097.

6. Schlichting, M., Richhariya, S., Herndon, N., Ma, D., Xin, J., Lenh, W., Abruzzi, K., and Rosbash, M. (2022). Dopamine and GPCR-mediated modulation of DN1 clock neurons gates the circadian timing of sleep. Proc Natl Acad Sci U S A 119, e2206066119. 10.1073/pnas.2206066119.

7. Zhang, Y., Liu, Y., Bilodeau-Wentworth, D., Hardin, P.E., and Emery, P. (2010). Light and temperature control the contribution of specific DN1 neurons to Drosophila circadian behavior. Curr Biol 20, 600–605. 10.1016/j.cub.2010.02.044.

8. Sun, L., Jiang, R.H., Ye, W.J., Rosbash, M., and Guo, F. (2022). Recurrent circadian circuitry regulates central brain activity to maintain sleep. Neuron 110, 2139–2154 e2135. 10.1016/j.neuron.2022.04.010.

9. Sato, K., and Yamamoto, D. (2023). Molecular and cellular origins of behavioral sex differences: a tiny little fly tells a lot. Front Mol Neurosci 16, 1284367. 10.3389/fnmol.2023.1284367.

10. Stockinger, P., Kvitsiani, D., Rotkopf, S., Tirián, L., and Dickson, B.J. (2005). Neural circuitry that governs male courtship behavior. Cell 121, 795–807. 10.1016/j.cell.2005.04.026.

11. Manoli, D.S., Foss, M., Villella, A., Taylor, B.J., Hall, J.C., and Baker, B.S. (2005). Male-specific fruitless specifies the neural substrates of Drosophila courtship behaviour. Nature 436, 395–400. 10.1038/nature03859.

12. Auer, T.O., and Benton, R. (2016). Sexual circuitry in Drosophila. Curr Opin Neurobiol 38, 18–26. 10.1016/j.conb.2016.01.004.

13. Yu, J.Y., Kanai, M.I., Demir, E., Jefferis, G.S., and Dickson, B.J. (2010). Cellular organization of the neural circuit that drives Drosophila courtship behavior. Curr Biol 20, 1602–1614. 10.1016/j.cub.2010.08.025.

14. Gummadova, J.O., Coutts, G.A., and Glossop, N.R. (2009). Analysis of the Drosophila Clock promoter reveals heterogeneity in expression between subgroups of central oscillator cells and identifies a novel enhancer region. J Biol Rhythms 24, 353–367. 10.1177/0748730409343890.

15. Talay, M., Richman, E.B., Snell, N.J., Hartmann, G.G., Fisher, J.D., Sorkac, A., Santoyo, J.F., Chou-Freed, C., Nair, N., Johnson, M., et al. (2017). Transsynaptic Mapping of Second-Order Taste Neurons in Flies by trans-Tango. Neuron 96, 783–795 e784. 10.1016/j.neuron.2017.10.011.

16. Nern, A., Pfeiffer, B.D., and Rubin, G.M. (2015). Optimized tools for multicolor stochastic labeling reveal diverse stereotyped cell arrangements in the fly visual system. Proc Natl Acad Sci U S A 112, E2967–2976. 10.1073/pnas.1506763112.

17. Fernandez, M.P., Berni, J., and Ceriani, M.F. (2008). Circadian remodeling of neuronal circuits involved in rhythmic behavior. PLoS Biol 6, e69. 10.1371/journal.pbio.0060069.

18. Dorkenwald, S., McKellar, C.E., Macrina, T., Kemnitz, N., Lee, K., Lu, R., Wu, J., Popovych, S., Mitchell, E., Nehoran, B., et al. (2022). FlyWire: online community for whole-brain connectomics. Nat Methods 19, 119–128. 10.1038/s41592-021-01330-0.

19. Nils Reinhard, A.F., Giulia Manoli, Emilia Derksen, Aika Saito, Gabriel Möller, Manabu Sekiguchi, Dirk Rieger, Charlotte Helfrich-Förster, Taishi Yoshii, Meet Zandawala (2024). Synaptic connectome of the Drosophila circadian clock. Biorxiv.

20. Dalton, J.E., Fear, J.M., Knott, S., Baker, B.S., McIntyre, L.M., and Arbeitman, M.N. (2013). Male-specific Fruitless isoforms have different regulatory roles conferred by distinct zinc finger DNA binding domains. BMC Genomics 14, 659. 10.1186/1471-2164-14-659.

21. von Philipsborn, A.C., Jorchel, S., Tirian, L., Demir, E., Morita, T., Stern, D.L., and Dickson, B.J. (2014). Cellular and behavioral functions of fruitless isoforms in Drosophila courtship. Curr Biol 24, 242–251. 10.1016/j.cub.2013.12.015.

22. Neville, M.C., Nojima, T., Ashley, E., Parker, D.J., Walker, J., Southall, T., Van de Sande, B., Marques, A.C., Fischer, B., Brand, A.H., et al. (2014). Male-specific fruitless isoforms target neurodevelopmental genes to specify a sexually dimorphic nervous system. Curr Biol 24, 229–241. 10.1016/j.cub.2013.11.035.

23. Anand, A., Villella, A., Ryner, L.C., Carlo, T., Goodwin, S.F., Song, H.J., Gailey, D.A., Morales, A., Hall, J.C., Baker, B.S., and Taylor, B.J. (2001). Molecular genetic dissection of the sex-specific and vital functions of the Drosophila melanogaster sex determination gene fruitless. Genetics 158, 1569–1595. 10.1093/genetics/158.4.1569.

24. Wohl, M., Ishii, K., and Asahina, K. (2020). Layered roles of fruitless isoforms in specification and function of male aggression-promoting neurons in Drosophila. Elife 9. 10.7554/eLife.52702.

25. Simpson, J.H. (2016). Rationally subdividing the fly nervous system with versatile expression reagents. J Neurogenet 30, 185–194. 10.1080/01677063.2016.1248761.

26. Kwon, Y., Kim, S.H., Ronderos, D.S., Lee, Y., Akitake, B., Woodward, O.M., Guggino, W.B., Smith, D.P., and Montell, C. (2010). TRPA1 Channel Is Required to Avoid the Naturally Occurring Insect Repellent Citronellal. Current Biology 20, 1672–1678. 10.1016/j.cub.2010.08.016.

27. Cichewicz, K., and Hirsh, J. (2018). ShinyR-DAM: a program analyzing Drosophila activity, sleep and circadian rhythms. Commun Biol 1, 25. 10.1038/s42003-018-0031-9.

28. Ozturk-Colak, A., Marygold, S.J., Antonazzo, G., Attrill, H., Goutte-Gattat, D., Jenkins, V.K., Matthews, B.B., Millburn, G., Dos Santos, G., Tabone, C.J., and FlyBase, C. (2024). FlyBase: updates to the Drosophila genes and genomes database. Genetics 227. 10.1093/genetics/iyad211.

29. Sanders, L.E., and Arbeitman, M.N. (2008). Doublesex establishes sexual dimorphism in the Drosophila central nervous system in an isoform-dependent manner by directing cell number. Dev Biol 320, 378–390. 10.1016/j.ydbio.2008.05.543.

